# Recurring functional interactions predict network architecture of interictal and ictal states in neocortical epilepsy

**DOI:** 10.1101/090662

**Authors:** Ankit N. Khambhati, Danielle S. Bassett, Brian S. Oommen, Stephanie H. Chen, Timothy H. Lucas, Kathryn A. Davis, Brian Litt

## Abstract

Human epilepsy patients suffer from spontaneous seizures, which originate in brain regions that also subserve normal function. Prior studies demonstrate focal, neocortical epilepsy is associated with dysfunction, several hours before seizures. How does the epileptic network perpetuate dysfunction during baseline periods? To address this question, we developed an unsupervised machine learning technique to disentangle patterns of functional interactions between brain regions, or subgraphs, from dynamic functional networks constructed from approximately 100 hours of intracranial recordings in each of 22 neocortical epilepsy patients. Using this approach, we found: (i) subgraphs from ictal (seizure) and interictal (baseline) epochs are topologically similar, (ii) interictal subgraph topology and dynamics can predict brain regions that generate seizures, and (iii) subgraphs undergo slower and more coordinated fluctuations during ictal epochs compared to interictal epochs. Our observations suggest that the epileptic network drives dysfunction by controlling dynamics of functional interactions between brain regions that generate seizures and those that underlie normal function.

## 1. Significance Statement

Localization-related epilepsy is a debilitating condition where seizures begin in dysfunctional brain regions, and is often resistant to medication. The challenge for treating patients is mapping dysfunction in brain networks that also subserve normal function several hours before seizures. Localizing brain regions that generate seizures is critical for improving seizure freedom rates following invasive surgery. We develop new methods to identify clusters of functionally interacting brain regions from approximately 100-hour intracranial, neocortical recordings per epilepsy patient. Our results indicate seizure-generating brain regions: (i) can be predicted before seizures and (ii) may kindle dysfunction through interactions with normal brain regions. These findings may have clinical implications for targeting specific brain regions to control seizures several hours before they occur.

## 2. Introduction

For approximately 60 million epilepsy patients, recurring, spontaneous seizures have a crippling impact on daily life. In approximately 26% of these patients, drivers of seizure activity have been linked to abnormal focal networks located in neocortical or mesial temporal structures [59]. To map dysfunction, epileptologists monitor continuous intracranial electrophysiology for biomarkers generated by the epileptic network, a set of interacting brain regions that are believed to initiate and spread seizure activity in the brain. To control seizures in medication-resistant individuals, clinical practitioners have traditionally prescribed resective surgery to remove brain tissue containing the epileptic network. More recently, epilepsy specialists are employing laser ablation and implantable devices to control dysfunction [60, 20, 49, 61, 47]. Novel neurotechnologies afford critical specificity in targeting brain circuits, but the key question for clinicians remains: “Which brain region(s) serve as the best target to control *this* patient’s seizures?”

Localizing epileptic brain regions based on abnormal electrophysiological biomarkers is a difficult problem, as etiology, seizure semiology, and frequency of events vary greatly between patients [39]. To reliably map the epileptic network, invasive monitoring lasts several days to weeks, and the length of the hospital stay increases the risks of infection, complications, and death. The extended monitoring period allows clinicians to describe a surgical target that accounts for variability in the seizure origin while minimizing expected impact on normal brain function. Recently, sampling error during limited monitoring time with intracranial electrodes has called into question the ability of traditional in-patient ictal recording to fully define the epileptic network [34]. This suggests that methods to map the epileptic network that do not rely on ictal recording may have significant advantages over current approaches. Critically, in localization-related epilepsy, brain regions that generate ictal (seizure) events are thought to be fundamentally altered in their structure and function, leading to the cognitive deficits observed during interictal (baseline) epochs [1, 40, 18, 26, 30]. These observations imply that brain circuits underlying cognitive functions are recruited by the epileptic network during interictal (baseline) states. However, when abnormal electrophysiology is not accompanied by discrete lesions evident on brain imaging, only about 40% of patients attain seizure freedom following resective surgery [21]. Modest outcomes associated with localization of abnormal electrophysiology suggests a fundamental gap in our understanding of how neurophysiologic biomarkers relate to pathophysiology in these patients.

A mechanistic understanding of seizure generation and evolution may be derived from spatial and temporal dynamics of the epileptic network [67, 28, 57, 54, 58, 70, 37, 29, 52, 66, 12, 23, 32, 31]. In this framework, network nodes are intracranial sensors measuring the electrocorticogram (ECoG) and network connections are time-varying statistical relationships between sensors [22, 27]. The degree of connectivity between brain regions is related to the synchronization of neural populations, a putative generator of dysfunction in epilepsy. Brain regions that are topologically central to the epileptic network tend to lie within [67, 28, 57, 58, 37, 29, 12, 32] and adjacent to [54, 70, 52, 66, 23] clinically-defined seizure-onset zones during interictal, preictal and ictal epochs [70, 65, 32]. In the context of this line of evidence, it is interesting to ask the question: “If network dysfunction persists over long time-scales, then (i) how does network topology drive brain dynamics differently during interictal and ictal epochs, and (ii) how might aberrant brain regions disrupt functional interactions underlying normal function?” Addressing these pressing questions about epileptic network physiology is crucial for targeting novel neurotechnology to dysfunctional brain circuits and minimizing impact on network structures involved in normal function.

In this work, we apply an unsupervised machine learning technique to examine how dynamic network architecture is differentially organized between ictal and interictal epochs. Our approach uncovers clusters of dynamic functional connections, or subgraphs, whose connection strengths undergo similar patterns of temporal variation, or expression, over several-day long ECoG recordings. Based on persistent network topology at the scale of ECoG [38], we first hypothesize that meso-scale functional networks form a repertoire of subgraphs, mapping out interactions between brain regions that recur through ictal and interictal epochs. The existence of recurring subgraphs might describe fundamental connections that guide network propagation of interictal epileptiform activity in trajectories similar to seizures [2, 41, 55, 69, 68, 35, 15, 31]. Second, we predict that functional subgraphs pinpoint connections specific to putative regions of seizure generation from normal functional connectivity within interictal epochs. Third, we hypothesize that functional subgraphs undergo slower, coordinated fluctuations in ictal epochs and faster, externally driven fluctuations in interictal epochs [32]. Our results support these hypotheses, demonstrating that functional subgraphs recur through ictal and interictal epochs, predict connectivity in the seizure-onset zone during interictal epochs, and differentiate ictal and interictal epochs on the basis of their time-varying expression.

## 3. Methods

### 3.1. Patient Data Sets

#### 3.1.1. Ethics Statement

All patients included in this study gave written informed consent in accordance with the Institutional Review Board of the University of Pennsylvania.

#### 3.1.2. Patient Demographics

#### 3.1.3. Electrophysiology Recordings

Twenty-two human patients (12 male and 10 female) undergoing surgical treatment for medically refractory epilepsy believed to be of neocortical origin underwent implantation of subdural electrodes to localize the seizure onset zone after non-invasive monitoring was indeterminate. De-identified patient data was retrieved from the online International Epilepsy Electrophysiology Portal (IEEG Portal) [64]. ECoG signals were recorded and digitized at either 512 Hz (Hospital of the University of Pennsylvania) or 500 Hz (Mayo Clinic) sampling rate. Surface electrode (Ad Tech Medical Instruments, Racine, WI) configurations, determined by a multidisciplinary team of neurologists and neurosurgeons, consisted of linear and two-dimensional arrays (2.3 mm diameter with 10 mm inter-contact spacing) and sampled the neocortex for epileptic foci (depth electrodes were first verified as being outside the seizure onset zone and subsequently discarded from this analysis). Signals were recorded using a referential montage with the reference electrode, chosen by the clinical team, distant to the site of seizure onset. Recordings spanned the duration of a patient’s stay in the epilepsy monitoring unit.

#### 3.1.4. Clinical Marking of the Seizure-Onset Zone

Seizure onset zone was marked on the Intracranial EEG (IEEG) according to standard clinical protocol in the Penn Epilepsy Center. Initial clinical markings are made on the IEEG the day of each seizure by the attending physician, always a board certified, staff epileptologist responsible for that inpatient’s care. Each week these IEEG markings are vetted in detail, and then finalized at surgical conference according to a consensus marking of 4 board certified epileptologists together. These markings on the IEEG are then related to other multi-modality testing, such as brain MRI, PET scan, neuropsychological testing, ictal SPECT scanning and magnetoenecephalographic findings to finalize surgical approach and planning. This process is standard of clinical care at National Association of Epilepsy Centers (NAEC) - certified Level-4 epilepsy centers in the United States.

#### 3.1.5. Description of Ictal and Interictal Epochs

Ictal epochs were identified by a team of neurologists as a part of routine clinical work and spanned the period between clinically-marked earliest electrographic change (EEC) [45] and termination. In this study, we disregarded sub-clinical seizures and only considered ictal epochs from clinical seizures that manifest seizure-related symptoms. Interictal epochs spanned 5 minutes in duration and were at least two hours removed from any ictal onset. We analyzed all possible interictal epochs from patient recordings.

### 3.2. Extracting Time-Varying Functional Networks

Signals from each 5-minute interictal epoch and each ictal epoch were divided into 1-second, non-overlapping, stationary time windows (**Fig 1A**) in accord with other studies [37] and subsequently pre-processed. In each time window, signals were re-referenced to the common average reference [62, 37] to account for variation in reference location across patients and to avoid broad field effects that may bias connectivity measurements erroneously in the positive direction. Each window was notch filtered at 60 Hz to remove line-noise, and low-pass and high-pass filtered at 120 Hz and 4 Hz, respectively, to account for noise, voltage drift, and *δ* frequency (0.5-4 Hz) contribution between time windows. To limit sources of volume conduction from introducing spurious connectivity, we pre-whiten signals in each window using a first-order autoregressive model to account for slow dynamics. Pre-whitening accomplishes two goals: (i) flattening of the signal power spectrum to enhance higher-frequency content that contains local neural population dynamics that is less affected by volume conduction, and (ii) decreasing the influence of independent node dynamics when computing correlation-based connectivity measurements [62, 10, 46, 3].

**Figure 1:**
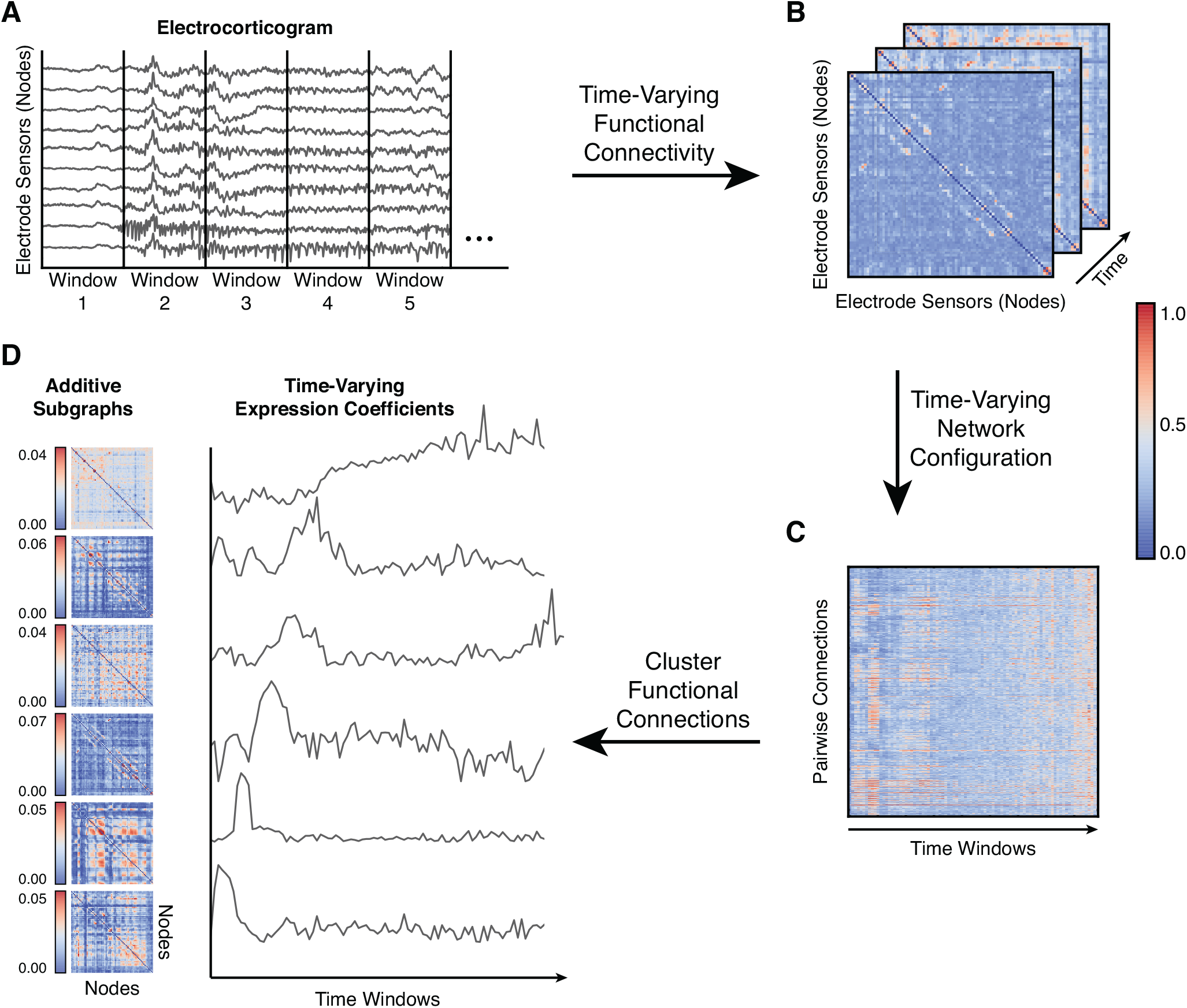
Clustering Functional Connections from Dynamic Epileptic Networks. (A) We identify ictal and interictal epochs from ECoG signals collected from patients with drug-resistant neocortical epilepsy implanted with intracranial electrodes. An ictal epoch is the period between seizure-onset – as characterized by the earliest electrographic change (EEC) [45] – and seizure termination. An interictal epoch is defined to be a continuous, 5 minute period at least 2 hours preceding or following seizure-onset. To measure time-varying functional networks, we divide each epoch into 1s time windows and estimate functional connectivity in each time window. In our model, each electrode sensor is a network node, and the weighted functional connectivity between sensors – interpreted as degree of synchrony – is represented as a network connection. (*B*) For each epoch, we estimated functional connectivity by applying a magnitude normalized cross-correlation between each pair of sensor time series in each time window. (*C*) For time-varying functional connectivity, we extract all pairwise connections between nodes and concatenate them over time windows to generate a time-varying network configuration matrix. (D) We apply *NMF* to the time-varying configuration matrix from each epoch, resulting in subgraphs that capture frequently repeating patterns of functional connections, and their expression over time.

Time-varying functional networks were formed by applying a normalized cross-correlation similarity function *ρ* between the time series of two sensors in the same time window using the formula, 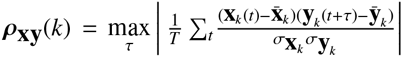, where **x** and **y** are signals from one of *N* sensors or network nodes, *k* is one of *K* non-overlapping, one-second time windows, *t* is one of *T* signal samples during the time window, *τ* = 1, 2, …, *T* is the time lag between signals, and *ρ* = 0 when node *x* is the same as node *y*. The *N* × *N* × *K* similarity matrix is also known as a time-varying adjacency matrix **A** (**Fig. 1B**). In our weighted network analysis, we retain and analyze all possible connection weights between nodes.

An alternate representation of the three-dimensional network adjacency matrix **A** is a two-dimensional network configuration matrix **Â**, which tabulates all *N* × *N* pairwise connection strengths across *K* time windows (**Fig. 1C**). Due to symmetry of **A**_*k*_, we unravel the upper triangle of **A**_*k*_, resulting in the weights of *N*(*N* − 1)/2 connections. Thus, **Â** has dimensions *N*(*N* − 1)/2 × *K*. We constructed a separate network configuration matrix for each ictal and interictal epoch.

### 3.3. Clustering Functional Connections into Subgraphs

To identify network subgraphs – sets of connections whose variation in strength cluster over time – we applied an unsupervised machine learning algorithm called non-negative matrix factorization (NMF) [42] to the network configuration matrix (**Fig. 1D**). This technique enabled us to pursue a parts-based decomposition of the time-varying network configuration matrix into subgraphs with time-varying expression coefficients [13]. Each subgraph is an additive component of the original network – weighted by its associated time-varying expression coefficient – and represents a pattern of functional interactions between brain regions. The NMF-based subgraph learning paradigm is a basis decomposition of a collection of dynamic graphs that separates co-varying network edges into subgraphs – or basis functions – with associated temporal coefficients – or basis weights. Unlike other graph clustering approaches that seek a hard partition of nodes and edges into clusters [6], the temporal coefficients provide a soft partition of the network edges, such that the original functional network of any time window can be reconstructed through a linear combination of all the subgraphs weighted by their associated temporal coefficient of that time window [43, 44, 13]. This implies that at a specific time window, subgraphs with a high temporal coefficient contribute their pattern of functional connections more than subgraphs with a low temporal coefficient.

Mathematically, NMF approximates **Â** by two low-rank, non-negative matrices, such that, **Â** ≈ **WH**, where **W** is the subgraph connectivity matrix (with dimensions *N*(*N* − 1)/2 × *m*), and **H** is the time-varying expression coefficients matrix (with dimensions *m*×*K*), and *m* is the optimized number of subgraphs learned. We applied NMF to the time-varying network configuration matrix using the alternating non-negative least squares with block-pivoting method with 100 iterations for fast and efficient factorization of large matrices [33]. We initialized **W** and **H** with non-negative weights drawn from a uniform random distribution on the interval [0, 1]. Due to the non-deterministic nature of this approach, we integrated subgraph estimates over multiple runs of the algorithm using *consensus clustering* – a general method of testing robustness and stability of clusters over many runs of one or more non-deterministic clustering algorithms [48]. Our adapted consensus clustering procedure [25, 24] entailed the following steps: (i) run the NMF algorithm *R* times per network configuration matrix, (ii) concatenate subgraph matrix **W** across *R* runs into an aggregate matrix with dimensions *N*(*N* − 1)/2 × *R* * *m*, and (iii) apply NMF to the aggregate matrix to determine a final set of subgraphs and expression coefficients.

In our study, we set *R* = 25 runs and separately repeated the consensus procedure for each epoch of each subject. We determined a subject-specific number of subgraphs *m* to learn across epochs by the following procedure: (i) randomly sample 50 epochs from the ictal and interictal pool, (ii) apply NMF for *m* = 2, 3, …, 20 subgraphs, independently for each epoch, (iii) compute the Frobenius error between **Â** and **WH** for each *m*, (iv) retain the value for *m* that occurs at the elbow of the resulting Frobenius error curve for each patient, and (v) find the optimum number of subgraphs 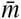 as the average *m* from the 50 epochs.

In sum, this subgraph learning procedure yielded *p* * 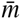 total subgraphs per patient, where *p* is the total number of ictal and interictal epochs.

#### 3.3.1. Generating Surrogate Subgraphs

An important mathematical property of subgraphs is that they form a basis set of the time-varying functional network from which they were derived. In other words, there exists a linear combination of an epoch’s subgraphs that reconstruct the original network, and any linear combination of the subgraphs forms a new subgraph that is still a basis of the original network. These properties allowed us to construct surrogate subgraphs with rewired network topology that maintain their basis functionality and preserve the empirically observed distribution of connection strengths.

We formed a set of surrogate subgraphs for each epoch by calculating a linear combination of the original subgraphs with weights pooled from a uniform random distribution on the interval [0, 1] (**Fig. 2A**). The size of the surrogate subgraph set remained equal to the size of the original subgraph set.

**Figure 2:**
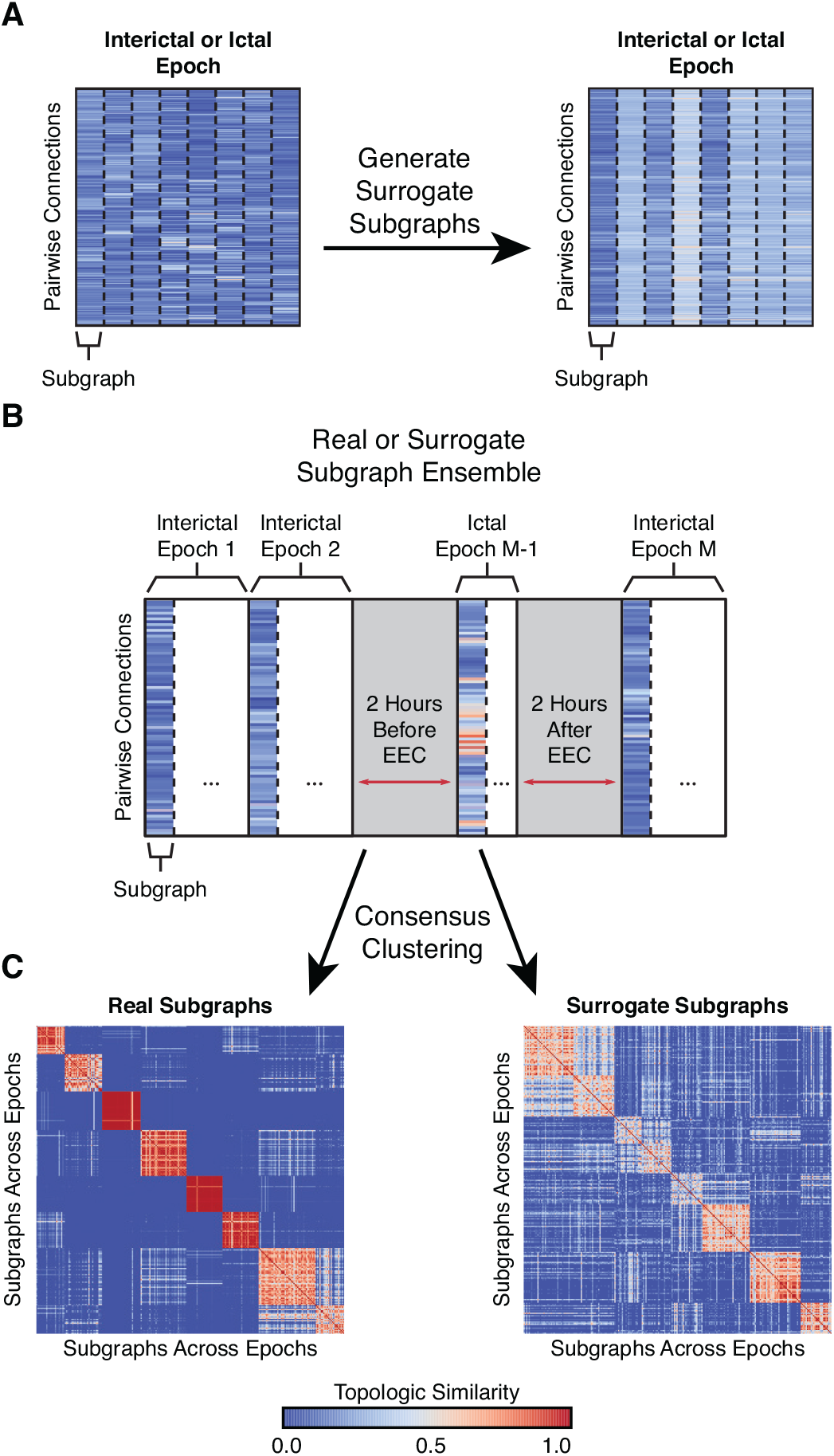
Clustering Subgraphs Based on Topological Similarity. (*A*) For the set of original subgraphs learned from an epoch of data (left), we generated an equally-sized set of surrogate subgraphs (right) by computing a weighted linear combination of the subgraphs with weights drawn from a uniform random distribution on the interval [0, 1]. The surrogate subgraphs have rewired network topology but maintain their functionality as a mathematical basis of the original network. (*B*) For each patient, we constructed a *subgraph ensemble matrix*, representing the *N*(*N* − 1)/2 functional connections for each subgraph from all interictal and ictal epochs. The ensemble matrix aggregates functional subgraphs expressed over approximately 100 hours of intracranial recording. We also constructed a patient-specific *surrogate ensemble matrix* by aggregating surrogate subgraphs across all epochs. (*C*) We quantified the topological similarity between all subgraphs in the ensemble matrix by applying a consensus NMF algorithm that tracks the number of times every pair of subgraphs is assigned to the same cluster over 100 runs of NMF. This procedure resulted in a co-clustering probability matrix representing the frequency with which subgraphs from ictal and interictal epochs are clustered together – a measure of similarity between the connectivity profiles of subgraph pairs. In the example, the co-clustering probability matrix of real subgraphs demonstrates less ambiguous similarity – matrix entries are near 0 or 1 – and greater clustering than surrogate subgraphs – matrix entries closer to 0.5.

### 3.4. Clustering Subgraph Ensembles Across Epochs

In this work, we sought to describe subgraph topology from the entirety of a patient’s data record by quantifying the similarity of subgraph connectivity profiles between interictal and ictal epochs. While several similarity and distance metrics are capable of comparing statistical features across observations in a data set (e.g. Pearson correlation, euclidean distance, cosine similarity), recent work has shown that a probabilistic measure of similarity derived from consensus clustering – by leveraging the non-deterministic property of the random initialization – may more accurately identify clusters in large data sets with many features [48]. To quantify topological similarity of subgraphs across all of a patient’s epochs, we again employed an NMF-based consensus clustering approach.

First, we compiled subgraphs across all of a patient’s epochs and constructed a subgraph ensemble matrix **E** (with dimensions 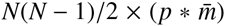) (**Fig. 2B**). To cluster the collection of *p* * 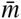 subgraphs, we applied multiple runs of NMF to **E,** such that, **E** ≈ **VG**, where **V** represents the subgraph for each cluster centroid (with dimensions *N*(*N* − 1)/2 × *j*) and **G** represents the likelihood cluster assignment for each subgraph (with dimensions 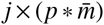), where *j* is the number of patient-wide clusters of subgraphs. After every NMF run, we retrieved the cluster assignment with maximum likelihood for each subgraph and counted the number of times each possible pair of subgraphs was assigned to the same cluster – and by extension the probability that any two subgraphs co-cluster [9, 25, 24]. These probabilities are tabulated in a symmetric co-clustering probability matrix **S** (with dimensions 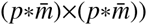 (**Fig. 2C**).

For every patient, we computed a co-clustering probability matrix **S** over 100 NMF runs for each number of subgraph clusters *j* = 2, 3, …, 20. To determine the optimum number of clusters *j*, we computed the Frobenius error between **E** and **VG** for each *j* and retained the value 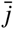 that occurs at the elbow of the resulting Frobenius error curve for each patient. Finally, we assigned each subgraph to its consensus cluster by applying one run of NMF, with 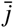 clusters, to **S.**

To generate a surrogate co-clustering probability matrix, we repeated our approach and replaced the original subgraphs in **E** with surrogate subgraphs and set the number of subgraphs *j* to the optimized number of subgraphs 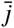 from the original ensemble clustering.

#### 3.4.1. Two-Dimensional Projection of Subgraph Similarity

To study the overall topological similarity between subgraphs, we employed a multi-dimensional scaling method [7] that projects each of the 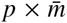 subgraphs as a data point in two-dimensional space and constrains the position of each data point a distance away from all other data points based on their relative similarities, as specified in **S**. In other words, more topologically similar (dissimilar) subgraphs are closer (further) in two-dimensional space (for example, see **Fig. 3A**). Formally, MDS assigns each subgraph a two-dimensional coordinate (xy) by minimizing the following stress function, Stress_**S**_ = 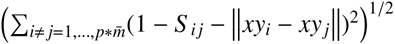, where **S** is the probabilistic subgraph co-clustering matrix, *i* and *j* are each different indices for one of 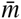 subgraphs of the *p* epochs. The MDS procedure assigns each subgraph a two-dimensional *xy* coordinate.

**Figure 3:**
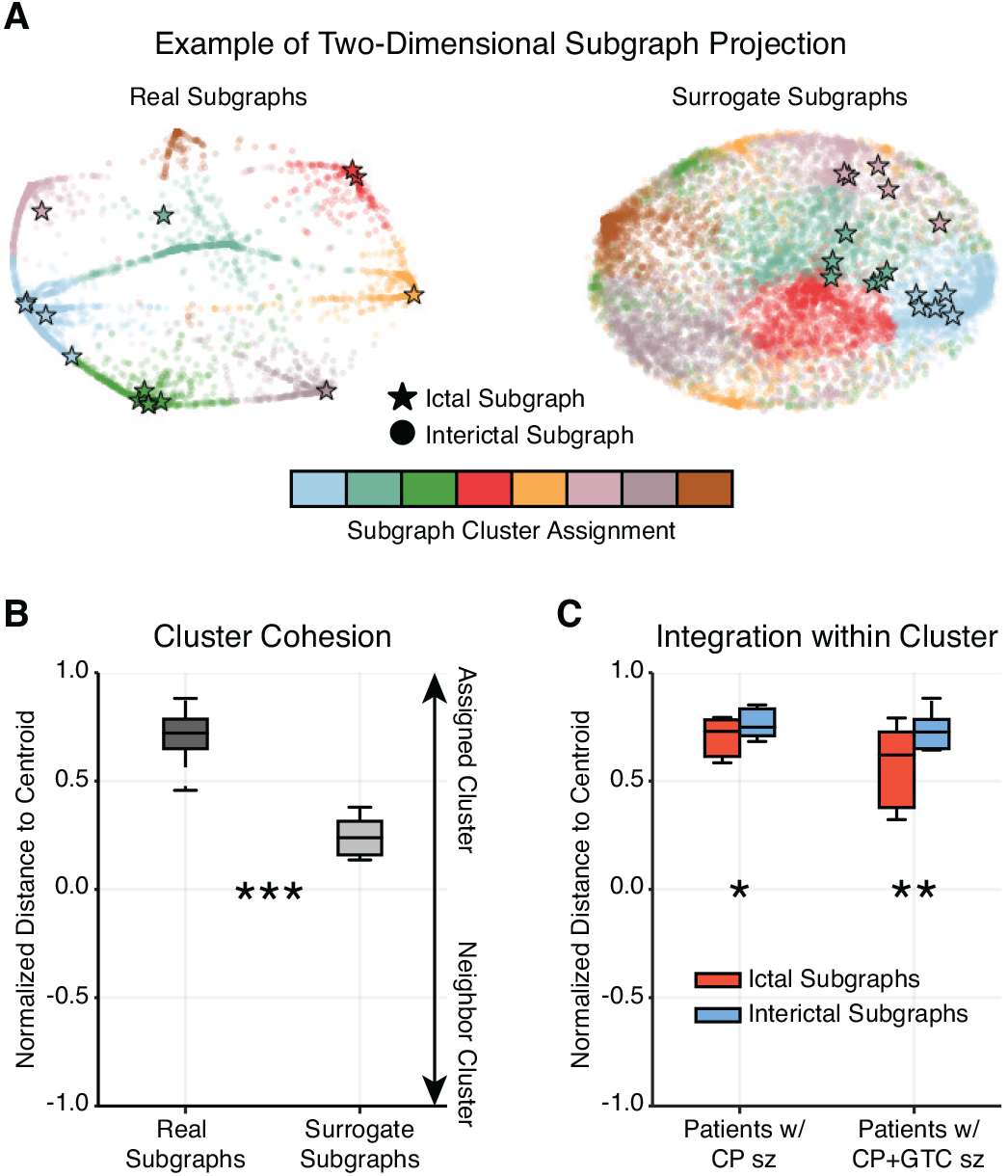
Ictal Subgraphs Are Recapitulated During Interictal Epochs. (**A**) Example two-dimensional projection of a patient’s subgraph co-clustering probability matrix. Each marker represents a subgraph from a single epoch and the distance between a subgraph pair indicates their topological similarity (i.e. closer subgraphs are more similar); circles represent interictal subgraphs and bolded stars represent ictal subgraphs; colors represent cluster assignment based on consensus clustering of the subgraph ensemble. The projections of real subgraphs (left) of the same cluster (color) tend to be closer to one another than to subgraphs of other clusters. In contrast, the projections of surrogate subgraphs from the same cluster tend to be as close to one another as surrogate subgraphs from other clusters. (**B**) Normalized, projected distance of a subgraph to its assigned cluster’s centroid – the mean geographical location of subgraphs in a cluster – relative to its neighboring cluster’s centroid (most proximal, non-assigned cluster centroid), averaged over all subgraphs of each patient (*N* = 22). Real subgraphs were significantly closer to their cluster centroid compared to surrogate subgraphs (paired *t*-test; *t*_21_ = 12.09, *p* < 7 × 10^−11^), suggesting the same set of brain regions functionally interact repeatedly over several hours. (**C**) Normalized, projected distance of ictal and interictal subgraphs to their cluster centroid, averaged over all ictal or interictal subgraphs of each patient with complex partial (CP) seizures (*N* = 8) and with secondarily-generalized complex partial (CP+GTC) seizures (*N* = 10). Both groups of patients expressed ictal subgraphs that were significantly further away from their cluster centroid than interictal subgraphs (paired *t*-test; CP: *t*_7_ = −3.29, *p* = 0.013; CP+GTC: *t*_9_ = −4.26, *p* = 0.002), suggesting ictal subgraphs may represent functional connections that lie at the transition between interictal subgraphs. (* *p* < 0.05, ** *p* < 0.01, *** *p* < 0.001; Bonferroni corrected)

Using the two-dimensional subgraph projection, we studied the proximity of a subgraph to its cluster centroid. Subgraphs closer to the centroid of their assigned cluster were considered more *integrated*, while subgraphs closer to the centroid of a non-assigned cluster (neighboring cluster) were considered more *promiscuous*. Formally, we computed a *normalized distance to centroid* measure by, Distance (*p, m, j*_assign_, *j*_neighbor_) = 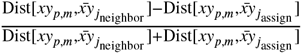, where *Dist* is the Euclidean distance function, *xy* are projected coordinates of the *m*^th^ subgraph of the *p*^th^ epoch, and 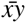 is the centroid coordinate of the assigned cluster for the subgraph *j*_assign_ or the centroid coordinate of the most proximal, non-assigned cluster *j*_neighbor_. Intuitively, a subgraph closer to the centroid of its assigned cluster than its neighboring cluster has normalized distance near +1, a subgraph closer to the centroid of its neighboring cluster than its assigned cluster has normalized distance near −1, and a subgraph equally distant to its own cluster centroid and neighboring cluster centroid has normalized distance of 0 (for example, see **Fig. 3B,C**).

### 3.5. Measures of Subgraph Topology and Dynahics

To quantify the topological and dynamic role of functional subgraphs in the epileptic network, we describe several measures based on the distributions of subgraph connectivity and expression coefficients.

To determine the degree to which a subgraph expressed functional connectivity in the seizure-onset zone, we computed a *SOZ sensitivity* measure of the relative strength of subgraph connectivity within the seizure-onset zone (SOZ) and outside the seizure-onset zone (OUT). Mathematically, the SOZ sensitivity is defined, SOZ Sensitivity(*p, m*) = 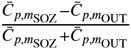, where 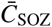 is the average subgraph connection strength of nodes within the SOZ and 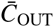 is the average subgraph connection strength of nodes outside the SOZ, of the *m*^th^ subgraph of the *p*^th^ epoch. The SOZ sensitivity ranges from +1, maximally sensitive to functional connections within the SOZ, to –1, maximally sensitive to functional connections outside the SOZ (for example, see **Fig. 4**). We also computed a surrogate distribution of SOZ sensitivity by randomly permuting the SOZ label across network nodes and re-computing SOZ sensitivity.

**Figure 4:**
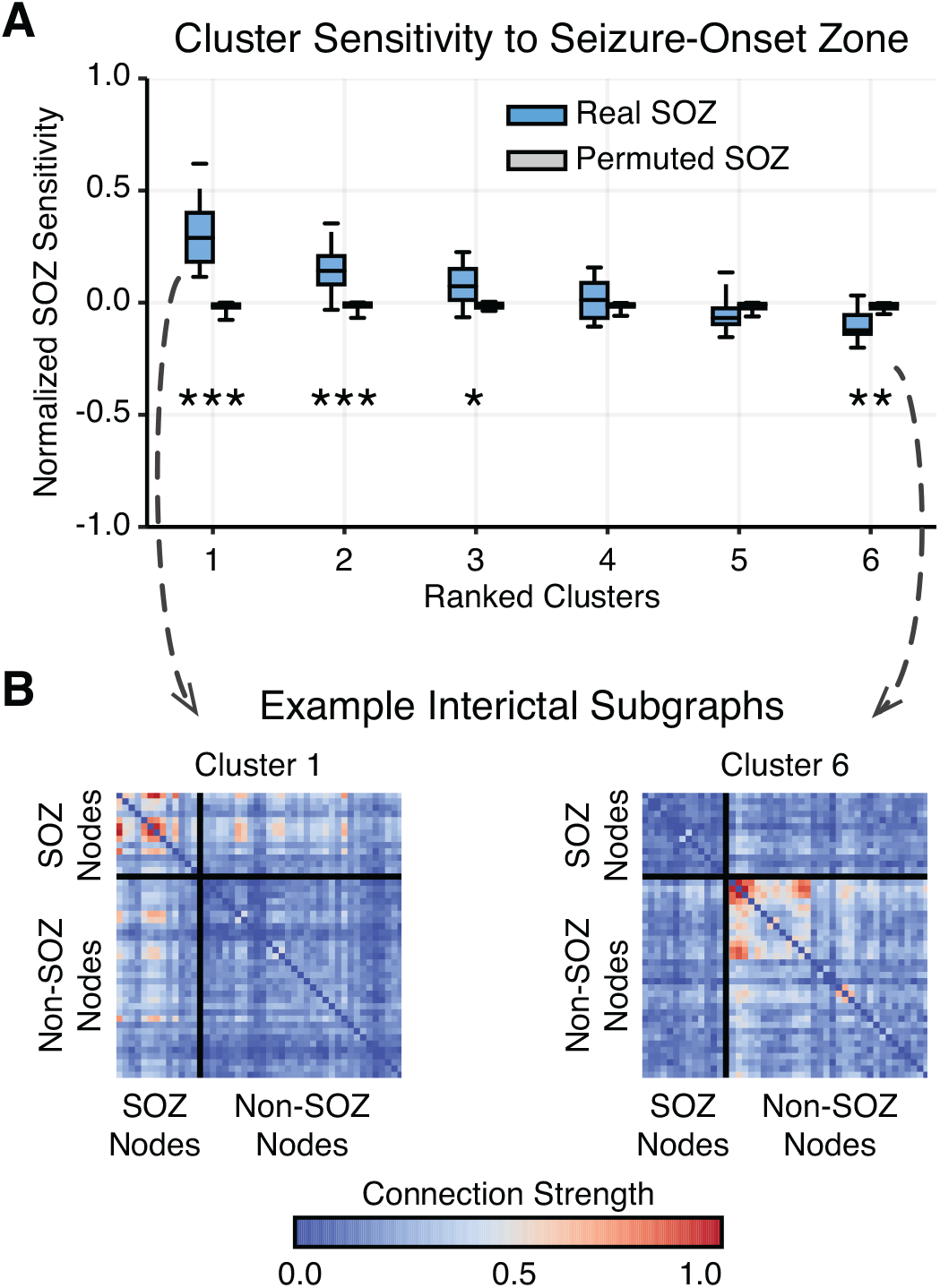
Interictal Subgraphs are Selectively Sensitive to the Seizure-Onset Zone. (**A**) Distribution of average SOZ sensitivity of subgraphs in each cluster, ranked in decreasing order, from each patient (*N* = 22). SOZ sensitivity of true SOZ labels in blue and of permuted SOZ labels in gray. We observed a significant effect of SOZ sensitivity for real SOZ labels compared to permuted SOZ labels for clusters 1, 2, 3, and 6 (* *p* < 0.05, ** *p* < 0.01, *** *p* < 0.001; Bonferroni corrected). These results demonstrate that functional interactions between brain regions are heterogeneously sensitive to dysfunction in the SOZ – depending on cluster-specific subgraph stereotypes. (**B**) Importantly, we observed that subgraphs of cluster 1 were significantly sensitive to connections within the SOZ, while subgraphs of cluster 6 were significantly sensitive to connections outside the SOZ. An example of subgraphs from cluster 1 (left) and cluster 6 (right) are shown here. Connections between SOZ nodes are shown in the top-left box, and connections between non-SOZ nodes are shown in the bottom-right box.

We compared subgraph dynamics between epochs by calculating the energy, skew, and power spectral density of subgraph expression coefficients. To compare subgraph expression between different epochs, we normalize each subgraph’s expression coefficients such that its maximum value is 1. The subgraph expression energy [13] quantifies the overall magnitude expression of the subgraph during an epoch (for example, see **Fig. 5C**) and was computed by, energy(p, m) = 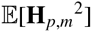, where **H** are the temporal coefficients of the *m*^th^ subgraph from the *p*^th^ epoch.

**Figure 5:**
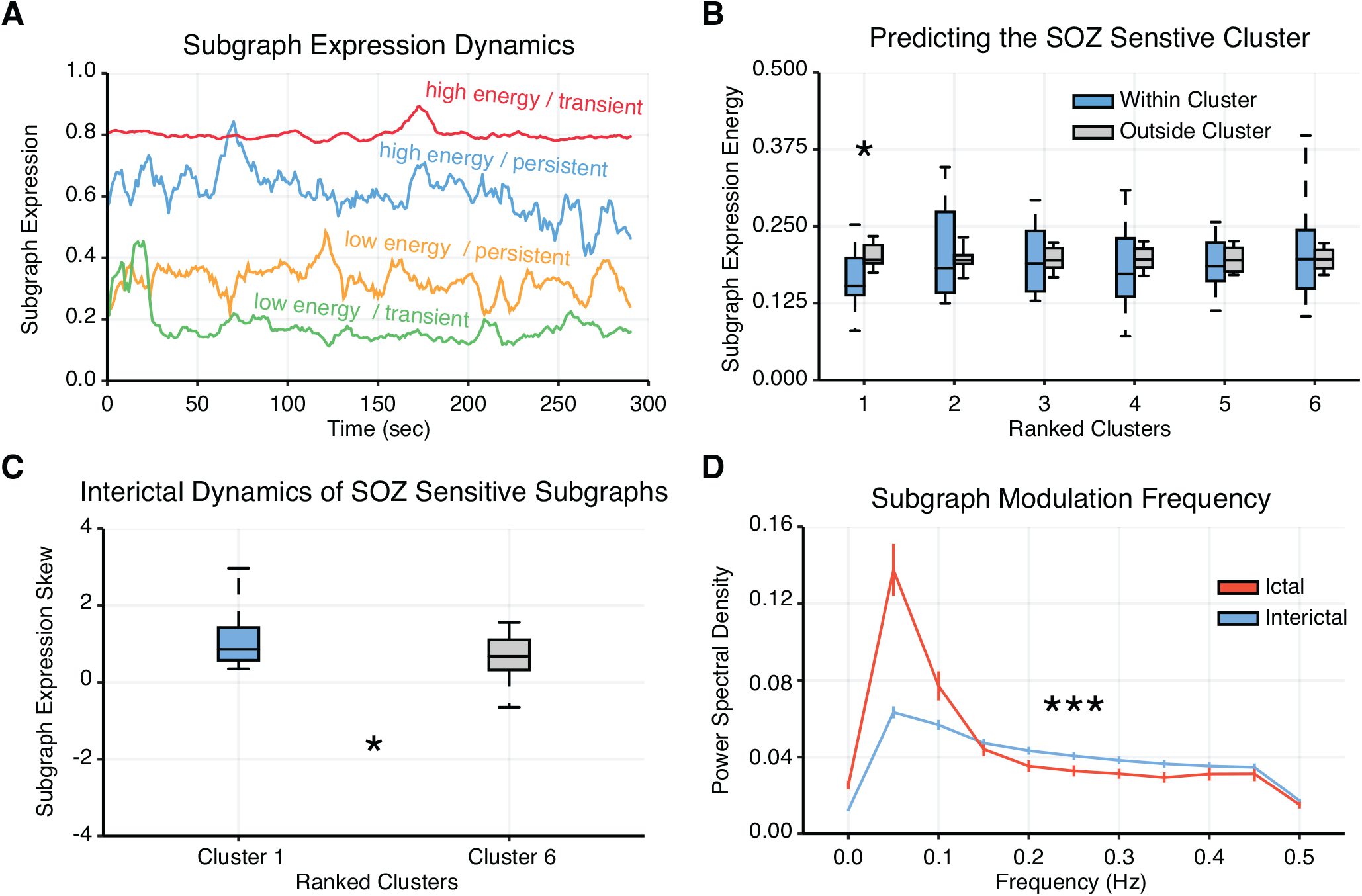
Expression Energy and Transience Differentiate Ictal and Interictal Epochs. (**A**) We computed subgraph expression energy – the overall activity of a subgraph – and subgraph expression skew – the temporal transience or persistence of a subgraph’s activity. Shown here are four examples of subgraph expression from a single patient – chosen by identifying subgraphs whose expression energy and expression skew were in the bottom and top third of all epochs – that demonstrate high energy and transience (red), high energy and persistence (blue), low energy and persistence (yellow), and low energy and transience (green). (**B**) Distribution of subgraph expression energy, averaged across interictal epochs of each cluster (ranked by SOZ sensitivity) for each patient (*N* = 22). For each cluster, we compared the distribution of expression energy for subgraphs of that cluster to expression energy for subgraphs of all other clusters and found significantly lower expression energy of subgraphs within cluster 1 – most sensitive to nodes in the SOZ – than outside cluster 1 (paired *t*-test; *t*_21_ = –3.21, *p* = 0.004; Bonferroni corrected for multiple comparisons). (**C**) Distribution of subgraph expression skew, averaged across interictal epochs of clusters 1 and 6 for each patient (*N* = 22). We observed subgraphs of cluster 1 – most sensitive to nodes in the SOZ – exhibited significantly greater skew, and therefore greater temporal transience, than subgraphs of cluster 6 – most sensitive to nodes outside the SOZ (paired *t*-test; *t*_21_ = 2.12, *p* = 0.04). These findings suggest that subgraphs with strongly connected SOZ nodes exhibit more transient, burst-like, dynamics than subgraphs with strongly connected non-SOZ nodes. (**D**) Power spectral density distribution of ictal and interictal subgraph expression, averaged over patients (*N* = 22). We observed a significant difference between ictal and interictal subgraph expression – ictal subgraphs modulate their expression at lower frequencies and interictal subgraphs modulate their expression at higher frequencies (Functional Data Analysis; *p* < 3 × 10^−5^). These findings suggest that subgraph expression is more gradual and coordinated during ictal epochs than interictal epochs.

The skew of a distribution of subgraph expression coefficients quantifies how transiently or persistently subgraphs are expressed [13]. Intuitively, transient subgraphs are expressed in brief, infrequent bursts – resulting in a heavy-tailed distribution of temporal coefficients (i.e., more small coefficients, and few large coefficients) – and persistent subgraphs are expressed evenly in time – resulting in a more normal distribution of temporal coefficients that fluctuate about the mean. The skew of the distribution of temporal coefficients for a subgraph distinguishes whether it is transiently (skew is greater than zero) or persistently (skew less than zero) expressed (for example, see **Fig. 5D**). The skew of the subgraph expression coefficients during an epoch, skew(*p, m*) 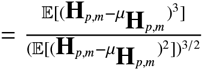, where **H** are the temporal coefficients of the *m*^th^ subgraph from the *p*^th^ epoch, and *µ*_*H*_ is the mean of the coefficients.

The power spectral density quantifies the modulation frequency of a subgraph’s expression [43] during an epoch and was computed using Welch’s Method with a sampling frequency of 1 Hz (each time window represents 1 second of functional connectivity) and a window size of 20 seconds (for example, see **Fig. 5D**).

## 4. Results

To disentangle functional subgraphs and their time-varying expression from epileptic brain, we retrieved ECoG recorded during ictal and interictal epochs from 22 patients undergoing routine pre-surgical evaluation of their neocortical epilepsy (see **Table 1** for patient-specific information) through the *International Epilepsy Electrophysiology Portal* (http://www.ieeg.org). We defined an ictal epoch as the period of ECoG signal between seizure-onset – as characterized by the earliest electrographic change (EEC) [45] – and seizure termination. Further, we defined an interictal epoch as a continuous 5 minute period of ECoG signal at least 2 hours preceding or following seizure-onset. We analyzed all possible interictal epochs, which amounted to *µ* = 106 ± 17 hours of ECoG signal per patient.

**Table 1:** Patient information. Patient data sets accessed through IEEG Portal (http://www.ieeg.org). Age at seizure-onset and at electrode implant surgery are noted. Location of seizure onset (lobe) and etiology are clinically-determined through medical history, imaging, and long-term invasive monitoring. Seizure types are SP (simple-partial), CP (complex-partial), CP+GTC (complex-partial with secondary generalization), or GA (generalized atonic). Counted seizures were recorded in the epilepsy monitoring unit. Interictal epochs were 5 minutes in duration and at least two hours away from any seizure. Clinical imaging analysis concludes L, Lesion; NL, non-lesion. Surgical outcome is reported by both Engel score and ILAE score (scale: I-IV/V, seizure freedom to no improvement; NR, no-resection; NF, no follow-up). M, male; F, female.

| Patient<br>(IEEG Portal) | Sex | Age<br>(Onset/<br>Surgery) | Seizure Onset | Etiology | Seizure<br>Type | Ictal<br>Epochs<br>(N) | Interictal<br>Epochs<br>(N) | Imaging | Outcome |
| --- | --- | --- | --- | --- | --- | --- | --- | --- | --- |
| HUP64_phaseII | M | 03/20 | Left frontal | Dysplasia | CP+GTC | 01 | 3228 | L | ENGEL-I |
| HUP65_phaseII | M | 02/36 | Right temporal | Auditory<br>reflex | CP+GTC | 03 | 2986 | N/A | ENGEL-I |
| HUP68_phaseII | F | 15/26 | Right temporal | Meningitis | CP,<br>CP+GTC | 05 | 3020 | NL | ENGEL-I |
| HUP70_phaseII | M | 10/32 | Left perirolandic | Cryptogenic | SP | 08 | 1079 | L | NR |
| HUP72_phaseII | F | 11/27 | Bilateral left | Mesial<br>temporal<br>sclerosis | CP+GTC | 01 | 2439 | L | NR |
| HUP73_phaseII | M | 11/39 | Anterior right<br>frontal | Meningitis | CP+GTC | 05 | 1071 | NL | ENGEL-I |
| HUP78_phaseII | M | 00/54 | Anterior left<br>temporal | Traumatic<br>injury | CP | 05 | 1719 | L | ENGEL-III |
| HUP79_phaseII | F | 11/39 | Occipital | Meningitis | CP | 01 | 1775 | L | NR |
| HUP86_phaseII | F | 18/25 | Left temporal | Cryptogenic | CP+GTC | 02 | 2612 | NL | ENGEL-II |
| HUP87_phaseII | M | 21/24 | Frontal | Meningitis | CP | 02 | 1201 | L | ENGEL-I |
| Study 004-2 | F | 14/27 | Right temporal<br>occipital | Unknown | CP+GTC | 01 | 638 | NL | ILAE-IV |
| Study 006 | M | 22/25 | Left frontal | Unknown | CP | 02 | 104 | NL | NR |
| Study 010 | F | 00/13 | Left frontal | Unknown | CP | 02 | 526 | L | NF |
| Study 011 | F | 10/34 | Right frontal | Unknown | CP,<br>CP+GTC | 02 | 283 | NL | NF |
| Study 016 | F | 05/36 | Right temporal<br>orbitofrontal | Unknown | CP+GTC | 03 | 669 | NL | ILAE-IV |
| Study 019 | F | 31/33 | Left temporal | Unknown | CP+GTC | 15 | 403 | NL | ILAE-V |
| Study 020 | M | 05/10 | Right frontal | Unknown | CP+GTC | 04 | 412 | NL | ILAE-IV |
| Study 023 | M | 01/16 | Left occipital | Unknown | CP | 04 | 208 | L | ILAE-I |
| Study 026 | M | 09/09 | Left frontal | Unknown | CP | 10 | 539 | NL | ILAE-I |
| Study 031 | M | 05/05 | Right frontal | Unknown | CP+GTC | 05 | 730 | NL | NF |
| Study 033 | M | 00/03 | Left frontal | Unknown | GA | 07 | 1321 | L | ILAE-V |
| Study 037 | F | 45/NR | Right temporal | Unknown | CP | 02 | 1087 | NL | NR |

**Table 2:** Statistical Table.

| Line | Data structure | Type of test | Power |
| --- | --- | --- | --- |
| a | Normal | paired $t$-test | 1 |
| b | Normal | paired $t$-test | 0.21 |
| c | Normal | paired $t$-test | 0.57 |
| d | Normal | paired $t$-test (Bonferroni corrected) | 1 |
| e | Normal | paired $t$-test (Bonferroni corrected) | 0.99 |
| f | Normal | paired $t$-test (Bonferroni corrected) | 0.97 |
| g | Normal | paired $t$-test (Bonferroni corrected) | 0.85 |
| h | Normal | paired $t$-test (Bonferroni corrected) | 0.10 |
| i | Normal | paired $t$-test (Bonferroni corrected) | 0.10 |
| j | Normal | paired $t$-test (Bonferroni corrected) | 0.55 |
| k | Normal | paired $t$-test | 0.58 |
| l | Nonparametric | Permutation test | $1 \times 10^6$ permutes |

For each epoch of each patient, we applied the following steps: (i) estimated weighted functional connectivity using a normalized cross-correlation metric and (ii) clustered patterns of frequently expressed functional connections from the network model by applying a machine learning technique called *non-negative matrix factorization (NMF)* to the time-varying network configuration matrix (see *Methods* for detailed procedure, and see Table 3 for number of subgraphs learned per epoch for each patient). This technique enabled us to pursue a parts-based decomposition of functional connections into subgraphs with time-varying expression coefficients [13]. Each subgraph is an additive component of the original network and represents a pattern of functional interactions between brain regions. Subgraphs are accompanied by time-varying expression coefficients, measuring the degree to which each subgraph is expressed at a given point in time.

**Table 3:** Subgraph Learning and Ensemble Clustering Table. Summary of number of ictal and interictal epochs, total number of epochs, optimized number of subgraphs learned per epoch, and optimized number of subgraph ensemble clusters for each patient.

| Patient<br>(IEEG Portal) | Electrode<br>Sensors<br>(N) | Electrode Configuration | Ictal<br>Epochs<br>(N) | Interictal<br>Epochs<br>(N) | Total<br>Epochs<br>($p$) | Subgraphs per<br>Epoch ($\bar{m}$) | Subgraph<br>Ensemble<br>Clusters ($\bar{j}$) |
| --- | --- | --- | --- | --- | --- | --- | --- |
| HUP64_phaseII | 88 | Grid: 8x8; Strip: 1x6 (4) | 01 | 3228 | 3229 | 8 | 8 |
| HUP65_phaseII | 80 | Grid: 8x8; Strip: 1x6 (3) | 03 | 2986 | 2989 | 8 | 9 |
| HUP68_phaseII | 79 | Grid: 8x8; Strip: 1x8 (2), 1x4 (2) | 05 | 3020 | 3025 | 8 | 7 |
| HUP70_phaseII | 78 | Grid: 8x8; Strip: 1x6, 1x4 (2) | 08 | 1079 | 1087 | 7 | 8 |
| HUP72_phaseII | 62 | Strip: 1x8 (3), 1x6 (5), 1x4 (2) | 01 | 2439 | 2440 | 8 | 9 |
| HUP73_phaseII | 56 | Strip: 1x8 (4), 1x6 (4) | 05 | 1071 | 1076 | 8 | 7 |
| HUP78_phaseII | 100 | Grid: 8x8; Strip: 1x6 (2), 1x4 (3);<br>Depth: 1x4 (3) | 05 | 1719 | 1724 | 6 | 8 |
| HUP79_phaseII | 84 | Grid: 6x8; Strip: 1x8, 1x6 (4), 1x4 | 01 | 1775 | 1776 | 8 | 8 |
| HUP86_phaseII | 118 | Grid: 8x8; Strip: 1x6 (5), 1x4 (4);<br>Depth: 1x4 (2) | 02 | 2612 | 2614 | 7 | 8 |
| HUP87_phaseII | 88 | Grid: 8x8; Strip: 1x4 (3); Depth: 1x4<br>(3) | 02 | 1201 | 1203 | 8 | 8 |
| Study 004-2 | 64 | Grid: 6x6; Strip: 1x4 (5); Depth: 1x4<br>(2) | 01 | 638 | 639 | 8 | 7 |
| Study 006 | 56 | Grid: 6x8; Strip: 1x8 | 02 | 104 | 106 | 8 | 8 |
| Study 010 | 56 | Grid: 6x8; Strip: 1x4 (2) | 02 | 526 | 528 | 8 | 10 |
| Study 011 | 84 | Grid: 6x8; Strip: 1x8 (2), 1x4 (5) | 02 | 283 | 285 | 7 | 7 |
| Study 016 | 64 | Grid: 4x6 (2); Strip: 1x4 (4) | 03 | 669 | 672 | 8 | 6 |
| Study 019 | 80 | Grid: 3x8, 6x6; Strip: 1x8 (2), 1x4 (3);<br>Depth: 1x4 (2) | 15 | 403 | 418 | 7 | 8 |
| Study 020 | 56 | Grid: 4x4, 4x6; Strip: 1x4 (4) | 04 | 412 | 416 | 8 | 9 |
| Study 023 | 92 | Grid: 8x8; Strip: 1x8, 1x4 (3); Depth:<br>1x4 (2) | 04 | 208 | 212 | 8 | 8 |
| Study 026 | 96 | Grid: 8x8; Strip: 1x8 (3), 1x4 (2) | 10 | 539 | 549 | 7 | 6 |
| Study 031 | 116 | Grid: 8x8, 4x6; Strip: 1x8 (2), 1x4 (3) | 05 | 730 | 735 | 7 | 7 |
| Study 033 | 124 | Grid: 8x8, 3x8; Strip: 1x8 (3), 1x4 (3) | 07 | 1321 | 1328 | 8 | 7 |
| Study 037 | 80 | Grid: 8x8; Strip: 1x8 (2) | 02 | 1087 | 1089 | 8 | 9 |

Importantly, our approach yields a collection of functional subgraphs over the long-term clinical recording. We studied the topology and dynamics of these learned subgraphs in greater detail to understand and pinpoint drivers of epileptic network dysfunction, interictally.

### 4.1. Ictal Network Architecture Emerges During Interictal Epochs

We first ask “Do subgraphs of interacting brain regions recur in their expression over the entire duration of a patient’s intracranial recordings?” We expected that if the same set of brain regions interact frequently, as described by a subgraph, then similar patterns of subgraph connectivity should emerge over the long-term recording. To test our hypothesis, we took the following probabilistic approach (**Fig. 2**): (i) constructed a *subgraph ensemble matrix* by aggregating functional connections over all subgraphs of a patient, (ii) quantified topological similarity between subgraphs by applying a consensus NMF algorithm to separate ensemble matrices for real and surrogate subgraphs, (iii) populated a real and a surrogate co-clustering probability matrix based on pairwise similarity of subgraphs from all epochs, and (iv) projected the co-clustering probability matrix on a two-dimensional Euclidean space using MDS. See Table 3 for number of subgraph ensemble clusters for each patient.

In the two-dimensional projection space, topologically similar subgraphs are geographically closer and topologically *dis*similar subgraphs are geographically farther from one another. We expected that interactions between brain regions prescribed by subgraphs within a cluster would be highly distinct from interactions between brain regions of other clusters. We visually confirmed this hypothesis in an sample patient, observing that geographically closer subgraphs were more likely assigned to the same cluster (**Fig. 3A**). In contrast, surrogate subgraphs, with randomized connectivity, of the same patient did not exhibit geographical clustering corresponding to the clustering assignment. To test whether clustering of topologically similar subgraphs is significantly greater in the true data than in the surrogate model, we quantified the degree of clustering by computing a normalized distance to centroid index for each subgraph that compares the Euclidean distance from a subgraph to its assigned cluster’s centroid and the same subgraph to its nearest neighboring cluster centroid (**Fig. 3B**). A cluster centroid is the mean two-dimensional, geographical location over all subgraphs in the cluster. Using a paired *t*-test, we found that the normalized distance to centroid, averaged over all subgraphs for each patient, was significantly greater for real subgraphs (*µ* = 0.71 ± 0.03) than surrogate subgraphs (*µ* = 0.24 ± 0.02; *t*_21_ = 12.09, *p* < 7 × 10^−11^; Table 2a). These results suggest that subgraphs assigned to the same cluster exhibit greater topological similarity than expected by chance. In other words, the functional architecture of meso-scale brain circuits is organized by recurring subgraphs of connectivity, in which the same sets of brain regions functionally interact, repeatedly, over several hours.

Based on our result of recurring functional subgraphs in epileptic brain, we next asked “Are ictal subgraphs topologically distinct from interictal subgraphs?” Visualizing the two-dimensional projection of the subgraph co-clustering probability matrix from an example patient (**Fig. 3A**), we observed several bridge-like extensions between subgraph clusters, representing putative transition graphs between clusters. We hypothesized that ictal subgraphs lie closer to the cluster periphery, at the junction of subgraph transitions, than interictal subgraphs. Moreover, we expected subgraphs of seizures that undergo more complex stages of spreading dynamics – secondarily-generalized, complex partial seizures (CP+GTC) – would be closer to these junctions (i.e. further from cluster centroid) than focal seizures whose dynamics minimally spread – complex partial seizures (CP). To test our hypothesis, we computed the normalized distance to centroid index, separately, for ictal and interictal subgraphs of each patient with CP seizures and with CP+GTC seizures (**Fig. 3C**). Using a paired *t*-test and Bonferroni multiple comparisons correction, we found: (i) for patients with CP seizures, ictal subgraphs were significantly more distant (*µ* = 0.70 ± 0.04) from their cluster centroid than interictal subgraphs (*µ* = 0.76 ± 0.03; *t*_7_ = −3.29, *p* = 0.013; Table 2b), and (ii) for patients with CP+GTC seizures, ictal subgraphs were significantly more distant (*µ* = 0.57 ± 0.06) from their cluster centroid than interictal subgraphs (*µ* = 0.71 ± 0.04; *t*_9_ = −4.26, *p* = 0.002; Table 2c). These results suggest that ictal subgraphs are less integrated within their clusters than interictal subgraphs and that ictal subgraphs of patients with CP+GTC seizures (*t*_9_ = −4.26) lie further from cluster centroid than ictal subgraphs of patients with CP seizures (*t*_7_ = −3.29). Importantly, ictal subgraphs are not topologically distinct from interictal subgraphs, and may, in fact, represent functional connections that lie at the transition between interictal subgraphs. Furthermore, seizures with complex patterns of spreading dynamics (CP+GTC) may express functional connections closer to junctions between subgraph clusters than seizures with more focal dynamics (CP).

### 4.2. Interictal Subgraphs Predict Seizure-Onset Regions

In the preceding analyses, we observed that: (i) ictal and interictal subgraphs that are more topologically similar are grouped in the same cluster and (ii) ictal subgraphs are topologically similar to interictal subgraphs and may capture transitions between clusters. If similar patterns of functional connectivity are expressed during ictal and interictal epochs, then we logically ask “Can interictal subgraphs predict which functional interactions drive seizure-onset?” To address this question, we compared interictal subgraph topology within and outside of clinically-defined seizure-onset brain regions. In accord with routine clinical evaluation of patients’ epilepsy, a team of neurologists successfully identified the sensors in the seizure-onset zone based on visual inspection of the intracranial recordings.

To determine the degree to which a subgraph expressed functional connectivity in the seizure-onset zone, we quantified the relative strength of brain regions within the seizure-onset zone (SOZ) and outside the seizure-onset zone (OUT) for each subgraph by computing the SOZ sensitivity measure.

We first asked, “Are all interictal subgraphs equally sensitive to connections in the SOZ, or are some interictal subgraphs more sensitive than others?” We hypothesized that connectivity in the SOZ would be expressed in a few interacting brain regions, rather than homogenously over many functional subgraphs. To test this hypothesis, we ranked subgraph clusters in decreasing order of their average SOZ sensitivity over inter-ictal subgraphs, for each patient. We expected cluster ranking to reveal potential hetereogeneity in the SOZ sensitivity of interictal subgraphs. Across the patient cohort, we generated a distribution of the average SOZ sensitivity for each of the top 6 ranked clusters – the minimum number of subgraph clusters identified for the 22 patients (**Fig. 4A**). Using a paired *t*-test and Bonferroni multiple comparisons correction, we compared the SOZ sensitivity distribution of each cluster to a null model in which brain regions within the SOZ are randomly permuted for every interictal subgraph. Compared to the null distribution, we found significantly greater SOZ sensitivity for cluster 1 (*µ* = 0.31 ± 0.04; *t*_21_ = 8.19, *p* < 2 × 10^−7^; Table 2d), cluster 2 (*µ* = 0.15 ± 0.03; *t*_21_ = 5.58, *p* < 3 × 10^−5^; Table 2e), and cluster 3 (*µ* = 0.08 ± 0.02; *t*_21_ = 3.75, *p* < 0.005; Table 2f), significantly lower SOZ sensitivity for cluster 6 (*µ* = −0.10 ± 0.02; *t*_21_ = −3.97, *p* < 0.001; Table 2g), and no significant difference for cluster 4 (*µ* = 0.01 ± 0.02; *t*_21_ = 1.86, *p* = 0.08; Table 2h) and cluster 5 (*µ* = −0.05 ± 0.02; *t*_21_ = −1.47, *p* = 0.16; Table 2i). These results suggest interictal subgraphs exhibit a heterogeneous sensitivity to brain regions within and outside the seizure-onset zone, with subgraphs in cluster 1 demonstrating the presence of network hubs localized to the SOZ and subgraphs in cluster 6 demonstrating the presence of network hubs localized outside the SOZ (**Fig. 4B**). Thus, interictal subgraphs express topological features that coincide with regions of dysfunction, for both strong connectivity and *dis*connectivity.

Next, we examine how these various functional subgraph topologies differentially behave in their pattern of time-varying expression – subgraph *dynamics*.

### 4.3. Temporal Dynamics Differentiate Subgraphs of Interictal and Ictal Epochs

We have presented evidence that ictal subgraphs are topologically similar to interictal subgraphs, and further that interictal subgraph topology can predict where seizures begin. Logically, we finally ask “If ictal and interictal subgraphs express similar network architecture, how is functional connectivity of the epileptic network differentially expressed between ictal and interictal epochs?" To answer this question, we analyzed the time-varying expression coefficients of each subgraph, which represent the degree to which a subgraph is expressed as a function of time. These coefficients are naturally provided by the NMF subgraph detection technique. From these data, we formulated two hypotheses: (i) that functional subgraphs express a variety of dynamical modes that predict subgraph topologies with heightened sensitivity for epileptic brain regions, and (ii) that expression of ictal subgraphs is modulated at slower time-scales than interictal subgraphs, supporting the notion that seizures are internally driven processes with coordinated dynamics.

To test our first hypothesis, we computed subgraph expression energy – a measure of overall dynamical activity – and subgraph expression skew – a measure of transient or persistent dynamics – and identified a sample of subgraphs that exhibit high/low energy and transient/persistence dynamics (**Fig. 5A**). We expected that expression energy would be predictably lower for interictal subgraphs with high SOZ sensitivity (cluster 1) compared to interictal subgraphs with lower SOZ sensitivity (cluster 2-6), accounting for more normal network dynamics during the seizure-free period. Using a paired *t*-test and Bonferroni multiple comparisons correction, we compared the distribution of expression energy, averaged over all interictal subgraph of each cluster, across patients to the distribution of expression energy, averaged over all interictal subgraphs outside that cluster, across patients (**Fig. 5B**). We found that interictal subgraphs of cluster 1 exhibit significantly lower expression energy (*µ* = 0.17 ± 0.01) than interictal subgraphs outside of cluster 1 (*µ* = 0.20 ± 0.004; *t*_21_ = –3.21, *p* = 0.004; Table 2j), suggesting that, indeed, subgraphs with high sensitivity to SOZ brain regions exhibit significantly attenuated activity during interictal epochs. Importantly, our results imply that expression energy is specific in its ability to predict the subgraph cluster that exhibits strong functional connections in the seizure-onset zone.

Next, we asked whether interictal subgraphs with pronounced connectivity in the SOZ (cluster 1) differ in their pattern of expression compared to interictal subgraphs with pronounced *dis*connectivity in the SOZ (cluster 6). We expected that subgraphs of cluster 1 may express their pattern of functional connections more intermittently, with greater transience, than subgraphs of cluster 6 (**Fig. 5C**). Using a paired *t*-test, we found that expression skew, averaged over all interictal subgraphs, across patients was greater for cluster 1 (*µ* = 1.25 ± 0.24) than cluster 6 (*µ* = 0.75 ± 0.20; *t*_21_ = 2.12, *p* = 0.04; Table 2k). These results suggest interictal subgraphs with high connectivity within the SOZ are expressed transiently, and interictal subgraphs with high connectivity outside the SOZ are expressed persistently.

Lastly, we tested our second hypothesis that topologically similar subgraphs differ in their expression frequency between ictal and interictal epochs. To test this hypothesis, we computed power spectral density (PSD) for each subgraph, averaged the PSD curves over all ictal or interictal subgraphs of each patient, and analyzed the resulting ictal and interictal PSD distribution (**Fig. 5A**). Using a statistical technique called *functional data analysis* (see [51] for technique, and [5] for illustrative application), we compared whether the area between ictal and interictal PSD curves were significantly different by comparing the true area to a null model in which ictal and interictal labels across subjects were permuted 1,000,000 times and the area between the curves was recomputed for each permutation. We found that the ictal and interictal PSD curves were significantly separated (area between curves = 0.014; *p* = 2.2 × 10^−5^; Table 21), suggesting ictal and interictal functional subgraph expression operates at different characteristic frequencies. Specifically, expression of ictal subgraphs modulates at slower frequencies and expression of interictal subgraphs modulates at higher frequencies – implying that similar patterns of functional connections of ictal and interictal subgraphs differ in their temporal dynamics. More generally, these results demonstrate that seizures mark a critical shift in network dynamics that is driven by slower and more coordinated expression of frequently interacting brain regions.

## 5. Discussion

In this work we asked, “Does interictal functional architecture of the epileptic brain perpetuate network dysfunction several hours between seizures?” To answer this question, we designed and applied a novel tool to disentangle subgraphs and their time-varying expression from dynamic functional connectivity. Our work supports the notion that ictal and interictal epochs traverse a similar set of functional subgraphs, but differ in the temporal pattern of subgraph expression – that is, subgraph *dynamics*.

### 5.1. Subgraphs Disentangle Regions of the Epileptic Network

A common notion in epilepsy is that dysfunctional cortical regions produce epileptiform activity, capable of generating seizures. However, network theorists posit that dysfunction may, in part, arise when neural activity between cortical regions hypersynchronize [63, 29]. Previous studies have identified discrete network states that describe shifts in global network topology, such as magnitude of functional connectivity [52, 12, 32]. However, these approaches are unable to pinpoint specific functional connections that drive changes in brain state across a seizure.

Building on prior work [17, 43, 44], in this study we disentangle functional networks into additive subgraphs, patterns of functional interactions between brain regions, that vary in expression over time. Logically, different subgraphs may be simultaneously or sequentially expressed to meet functional demand [4, 16, 11, 53, 13]. Our results demonstrate that the dynamic epileptic network expresses functional subgraphs that recur during ictal and interictal epochs. It is intuitively plausible that the epileptic network is actually composed of a small set of subgraphs that underlie normal function during interictal epochs, but are co-opted to support seizure dynamics during ictal epochs [56, 36, 50, 35, 32]. Such a theory is corroborated by our finding that subgraphs of ictal epochs are more likely to lie at the transition between clusters representing different gross topological architecture and exhibit slower and more coordinated dynamics than during interictal epochs.

Importantly, the geography of the subgraph projection space points to a core-periphery organization [8] of ictal and interictal subgraphs – in which more densely clustered interictal subgraphs form a core set of highly similar topologies and more loosely clustered ictal subgraphs form a network periphery of more variable topologies. The existence of core-periphery organization in dynamical brain networks related to language [19, 14] and learning [6], supports the idea that temporally-variable network architectures help navigate different cognitive states. In the epileptic network, ictal subgraphs of the cluster periphery may be more likely to facilitate dynamical transitions between clusters of different subgraph topologies than interictal subgraphs. Furthermore, our finding that subgraphs of seizures with pronounced spatial spread (CP+GTC) lie closer to their cluster periphery than focal seizures (CP) may contribute to global properties of network topology that have been used to predict seizure type in prior work [31]. Neurophysiologically, the epileptic network demonstrates a weakened regulatory, push-pull control in constraining CP+GTC seizures [31] and might contribute to the ability of CP+GTC subgraphs to more flexibly transition between subgraph clusters than CP subgraphs.

### 5.2. Predicting Seizure Origin in the Network

We observed that functional interactions specific to the seizure-onset zone are highly predicted by the magnitude of functional connectivity and cluster assignments of topologically similar, interictal subgraphs. Our results agree with prior studies demonstrating increased network connectivity in seizure onset regions during interictal epochs [65, 35]. Our finding that topologically similar subgraphs form clusters over the long data record suggests that the pattern of functional interactions is critical to differentiate regions that drive seizure onset from the surrounding network.

Importantly, our results demonstrate that the site of seizure origin in the epileptic network exhibits dysfunction that recurs transiently over long periods of time. Furthermore, our novel subgraph clustering approach reliably pinpoints this target several hours before seizures occur and reveals that the region is overall more "silent" or dormant relative to more normal brain networks. However, we witnessed that these dysfunctioned and attenuated subgraphs can transiently disrupt functional interactions underlying more normal and persistent brain processes. Prior work has shown that focal, left-sided epileptiform activity is associated with decreased short-term verbal memory and focal, right-sided epileptiform activity is associated with decreased short-term memory in non-verbal or spatial tasks [1, 26]. Further studies demonstrate that seizures originating in the temporal lobe result in decreased cognitive performance on tasks often associated with activation of frontal and prefrontal lobe, such as performance IQ, verbal IQ, and word list learning [30], suggesting that cognitive functions are impacted over long distances through network interactions. The approach we developed can be used to study pressing questions regarding secondary deficits caused by interactions between epileptic and normal brain regions.

### 5.3. Methodological Limitations and Extensions

The first important clinical consideration related to this work is the sampling error inherent in any intracranial implantation procedure. Any of the techniques used to map epileptic brain usually yield incomplete representations of the epileptic network. As a consequence, the subgraphs we measured may represent just a portion of more distributed functional circuits that extend further throughout the brain.

Secondly, our methods of predicting epileptic network architecture from interictal epochs relies on accurate delineation of seizure-onset regions. Because of sampling error and variability in clinical decision-making, the seizure-onset region may be under- or over-sampled. However, the goodness-of-fit of our statistical model in predicting seizure-onset regions based on functional connectivity suggests that our model reasonably agrees with a consensus definition of the seizure-onset zone formed by a team of practicing neurologists.

### 5.4. Clinical Impact

Mapping architecture of the epileptic network presents significant challenges for clinicians. In patients with neocortical epilepsy, we showed that functional network topology is highly similar between ictal and interictal epochs. These findings are relevant for (i) optimizing treatment strategies to reduce dysfunction and preserve normal function, and (ii) reducing morbidity and mortality associated with extended duration of invasive intracranial electrode implantation, which according to recent studies, may actually require months of outpatient intracranial recording with implantable devices [34]. By predicting seizure-onset regions from interictal epochs, clinical monitoring may be shortened, or potentially even conducted intraoperatively. In this setting, one might imagine epilepsy surgery or device placement taking place in one procedure, relying on interictal brain network mapping, delivered similarly to ablations performed by cardiac electrophysiologists. Furthermore, our finding that complex patterns of functional connectivity correlate with sources of dysfunction supports the use of novel interventional strategies, such as laser ablation or implantable devices, to affect functional circuits at finer spatial scales than is currently possible with large resective surgery.

## Acknowledgments

AK and BL acknowledge support from the National Institutes of Health through awards #R01-NS063039, #1U24 NS 63930-01A1, the Citizens United for Research in Epilepsy (CURE) through Julie’s Hope Award, and the Mirowski Foundation. DSB acknowledge support from the John D. and Catherine T. MacArthur Foundation, the Alfred P. Sloan Foundation, the Army Research Laboratory and the Army Research Office through contract numbers W911NF-10-2-0022 and W911NF-14-1-0679, the National Institute of Mental Health (2-R01-DC-009209-11), the National Institute of Child Health and Human Development (1R01HD086888-01), the Office of Naval Research, and the National Science Foundation (BCS-1441502, BCS-1430087, and PHY-1554488). The content is solely the responsibility of the authors and does not necessarily represent the official views of any of the funding agencies.

